# An exploratory study of gastrointestinal redox biomarkers in the presymptomatic and symptomatic Tg2576 mouse model of familial Alzheimer’s disease – phenotypic correlates and the effects of chronic oral D-galactose

**DOI:** 10.1101/2023.06.03.542513

**Authors:** Jan Homolak, Ana Babic Perhoc, Ana Knezovic, Jelena Osmanovic Barilar, Davor Virag, Melita Salkovic-Petrisic

## Abstract

The gut might play an important role in the etiopathogenesis of Alzheimer’s disease (AD) as gastrointestinal alterations often precede the development of neuropathological changes in the brain and correlate with disease progression in animal models. The gut has an immense capacity to generate free radicals whose role in the etiopathogenesis of AD is well known; however, it remains to be clarified whether gastrointestinal redox homeostasis is associated with the development of AD. The aim was to i) examine gastrointestinal redox homeostasis in the presymptomatic and symptomatic Tg2576 mouse model of AD; ii) investigate the effects of chronic oral D-galactose previously shown to alleviate cognitive deficits and metabolic changes in animal models of AD; iii) investigate the association between gastrointestinal redox biomarkers and behavioral alterations in Tg2576 mice. Presymptomatic Tg2576 have a heightened gastrointestinal electrophilic tone reflected in increased lipid peroxidation and activity of Mn/Fe-SOD. Chronic oral D-galactose treatment was associated with detrimental effects on redox homeostasis only in the wild-type controls. In the symptomatic stage, Tg2576 mice demonstrate compensated redox disbalance characterized by normalized lipid peroxidation and increased hydrogen peroxide dissociation capacity but diminished total antioxidant reserve alleviated with chronic oral D-galactose treatment. Conversely, D-galactose reduced antioxidant capacity and increased lipid peroxidation in the controls. Total antioxidant capacity was associated with greater spatial memory, while other biomarkers had a complex relationship with exploration, nesting, and grooming. Gut redox homeostasis might be involved in the development and progression of AD pathophysiology and should be further explored in this context.

## Introduction

Accumulating evidence suggests gut might play an important role in the etiopathogenesis and progression of Alzheimer’s disease (AD)^1,2^. Gastrointestinal (GI) alterations have been reported in both transgenic^3–7^ and non-transgenic animal models of AD^8–11^. Importantly, GI alterations often precede neuropathological changes in the brain and correlate with disease progression^4,6^, suggesting pathophysiological processes in the gut might contribute to dyshomeostasis in the central nervous system (CNS) in the early stages of neurodegeneration. Honarpisheh et al. reported that GI dysfunction takes place before the onset of cognitive symptoms and the accumulation of cerebral amyloid-β (Aβ) in the (6-month-old) Tg2576 mouse model of familial AD^4^. Similarly, pathophysiological alterations in the GI tract have been observed in the presymptomatic stage in other animal models of familial AD (e.g. TgCRND8^6^ and APP/PS1^3^). Mechanisms by which gut dyshomeostasis might incite neurodegeneration are still not clear; however, current working models suggest dysfunction of the GI barrier might promote chronic inflammation and metabolic dyshomeostasis associated with microglial activation and brain insulin resistance as key etiopathogenetic clusters of AD^12–16^. GI redox dyshomeostasis might be another mechanism by which gut dysfunction impels neurodegenerative processes: i) oxidative stress is closely related to most molecular mechanisms driving AD^17,18^; ii) the gut has an immense capacity to generate free radicals with detrimental health consequences^19–21^ and 40% of body energy expenditure is required for the maintenance of the GI barrier which is constantly and inevitably exposed to xenobiotics and microorganisms^22^; iii) maintenance of gut redox homeostasis and the GI barrier are mutually intertwined physiological processes^21,23,24^. So far, redox homeostasis has only been examined in the presymptomatic APP/PS1 mouse model of familial AD demonstrating a reduced gut antioxidant capacity and diminished expression of proteins involved in the maintenance and utilization of glutathione (GSH)^3^. Here, we aimed to examine redox homeostasis in the GI tract of the presymptomatic and symptomatic Tg2576 mice, one of the most widely used transgenic models of AD. Our second aim was to assess the effects of chronic oral D-galactose treatment in the presymptomatic and symptomatic Tg2576 and their age-matched controls. Chronic oral D-galactose treatment initiated in the early post-induction period has the potential to prevent and alleviate cognitive dysfunction in the rat model of sporadic AD^25,26^ and recent evidence indicates that oral D-galactose can modulate redox homeostasis in the brain^27^ and exert favorable effects on redox signaling in the gut^28^. Finally, considering the importance of the gut-brain axis in regulating behavior^29–31^, we aimed to examine phenotypic correlates GI redox biomarkers using data from a behavioral assessment conducted in the same cohort of Tg2576 mice by Babic Perhoc et al.^32^.

## Results

Presymptomatic Tg2576 had increased activity of gut Mn-SOD (−14% in δTHB absorbance (inversely proportional to Mn-SOD activity); p = 0.004). Increased activity of Mn-SOD was associated with increased lipid peroxidation (+59%) which did not reach the predetermined statistical significance threshold due to large variability. Chronic oral D-galactose treatment (200 mg/kg; *ad libitum*) was not associated with pronounced changes in gastrointestinal redox biomarkers. Total antioxidant capacity was unaltered. In the wild-type controls, chronic oral D-galactose treatment was associated with decreased H_2_O_2_ dissociation capacity (−25%) and increased baseline H_2_O_2_ (+26%), possibly due to suppressed activity of catalase (−26%; p = 0.08). D-galactose treatment also increased the activity of SODs, namely the mitochondrial Mn-SOD (−13% δTHB absorbance) (Table 1; Figure 1) and increased lipid peroxidation (+17%). Interestingly, the observed effect was largely absent in the presymptomatic Tg2576. There was no change in total H_2_O_2_ dissociation capacity, the activity of catalase and SOD, nor the concentration of lipid peroxidation end products (Table 1; Figure 1).

**Figure 1.**
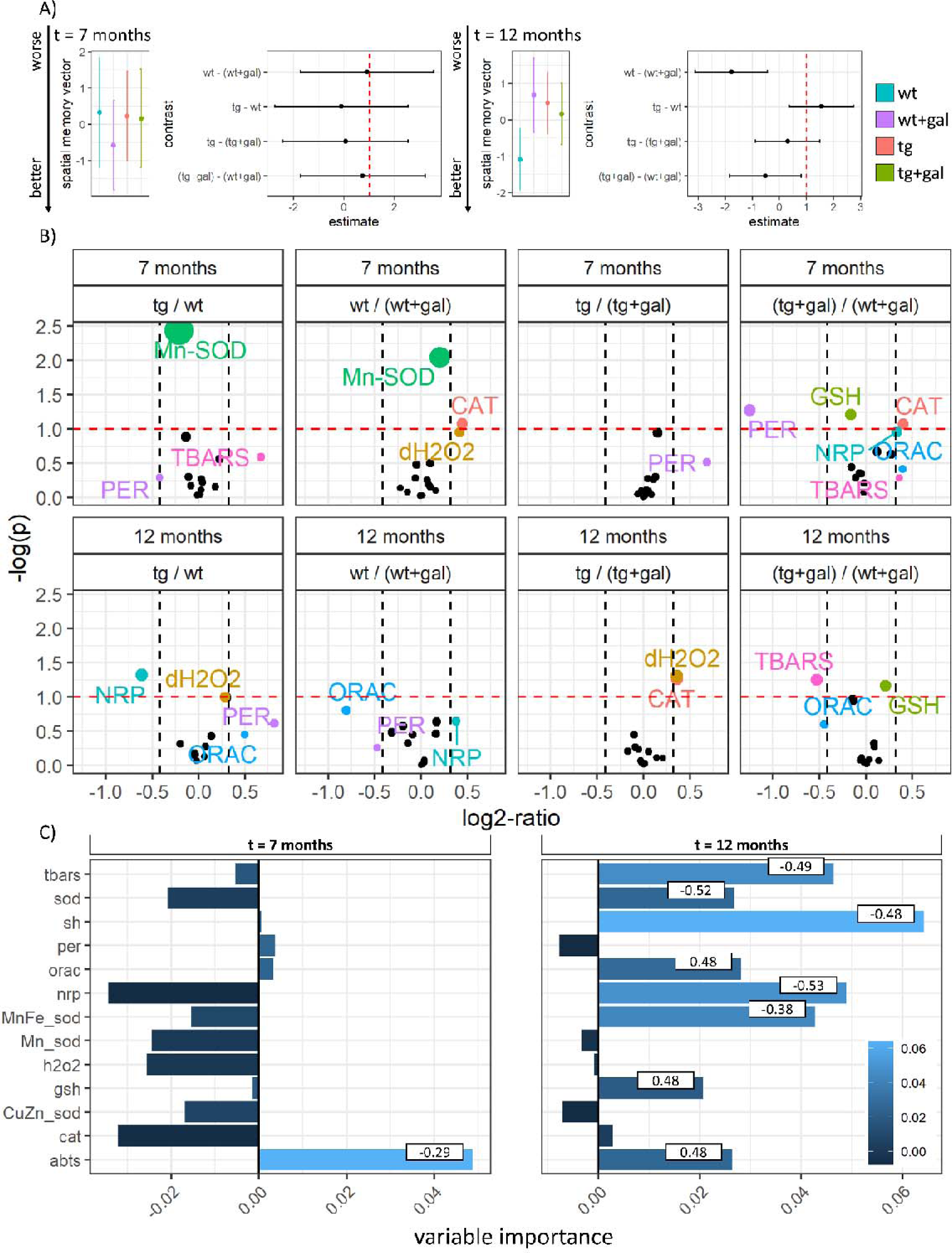
Spatial memory and gastrointestinal redox biomarkers in the presymptomatic (7- month-old) and symptomatic (12-month-old) Tg2576 mice after chronic oral D-galactose treatment (200 mg/kg). **A)** Model output depicting spatial memory vector (SMV) point estimates with 95% confidence intervals (CI) in the presymptomatic (left) and symptomatic (right) Tg2576 mice and respective wild-type controls treated with vehicle (wt; tg) or oral D-galactose (wt+gal; tg+gal). Larger values are associated with worse performance in Morris Water Maze spatial memory test. Group differences are presented as contrasts (point estimates and 95% CI of differences between group least square means). **B)** Volcano plot demonstrating group comparisons (ratiometric contrasts) with log_2_ of ratios on the X-axis and -log_10_ of p-values on the Y-axis. The exploratory threshold for -log_10_(p) is presented as a horizontal red dotted line and thresholds of the effect size (set at 25%) are denoted as vertical black dotted lines. **C)** Variable importance maps calculated based on the permutation-induced mean decrease in accuracy derived from conditional inference-based unbiased classification random forests with SMV defined as the response variable. Bars indicate permutation importance for cforest models and numbers indicate correlation coefficients. abts – difference in absorbance of 2,2′-azino-bis(3-ethylbenzothiazoline-6-sulfonic acid); cat – catalase activity; Cu/Zn-sod – the activity of cytoplasmic Cu/Zn-superoxide dismutase; dh2o2 – total H_2_O_2_ dissociation capacity; gsh – glutathione; h2o2 – H_2_O_2_; Mn-sod – Mn-superoxide dismutase; Mn/Fe-sod – Mn- and Fe-superoxide dismutase; nrp – nitrocellulose redox permanganometry (antioxidant capacity biomarker); orac – oxygen radical absorbance capacity; per – residual activity of peroxidases; sh – protein thiol residues; sod – total superoxide dismutase activity; tbars – thiobarbituric acid reactive substances.

**Table 1.**
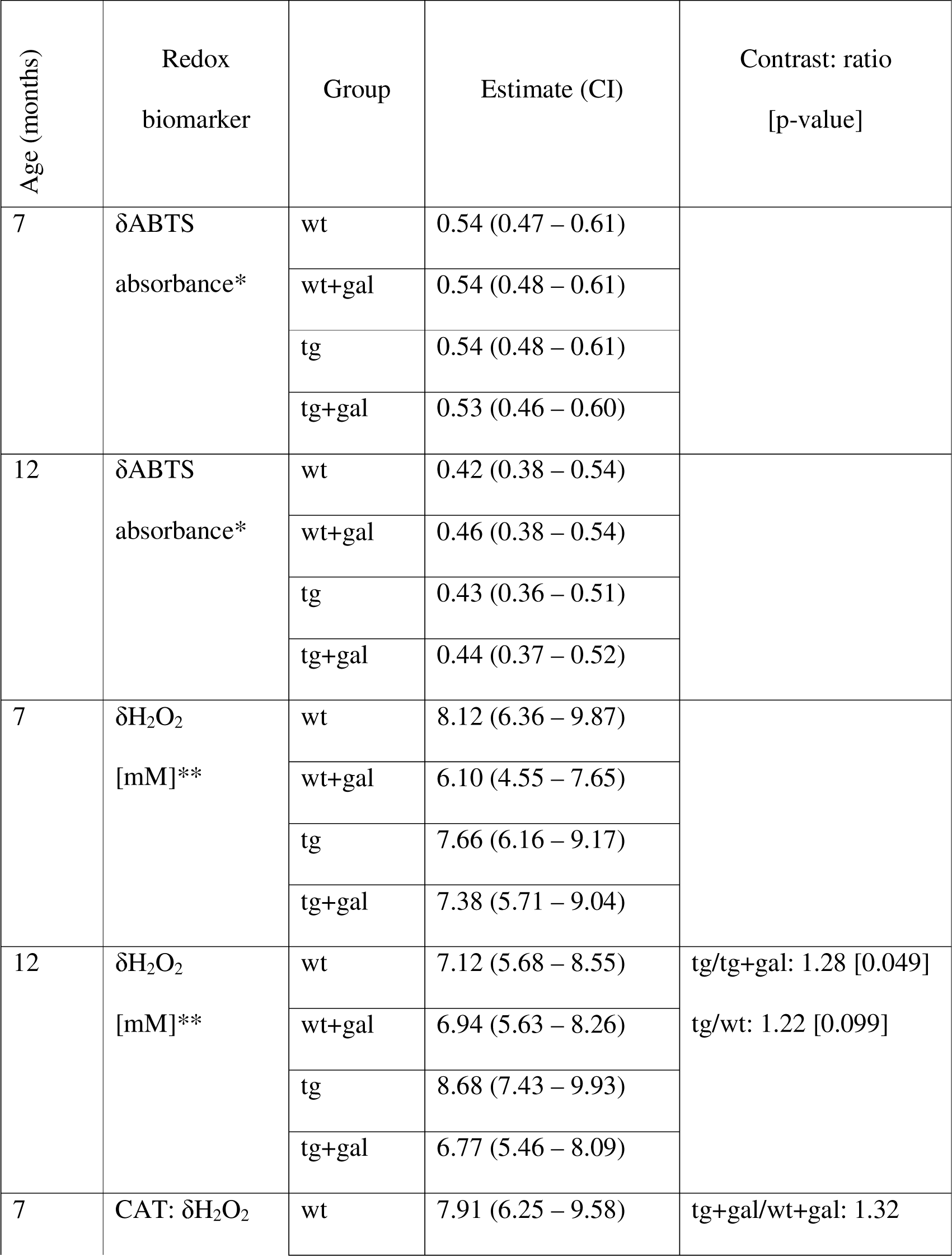
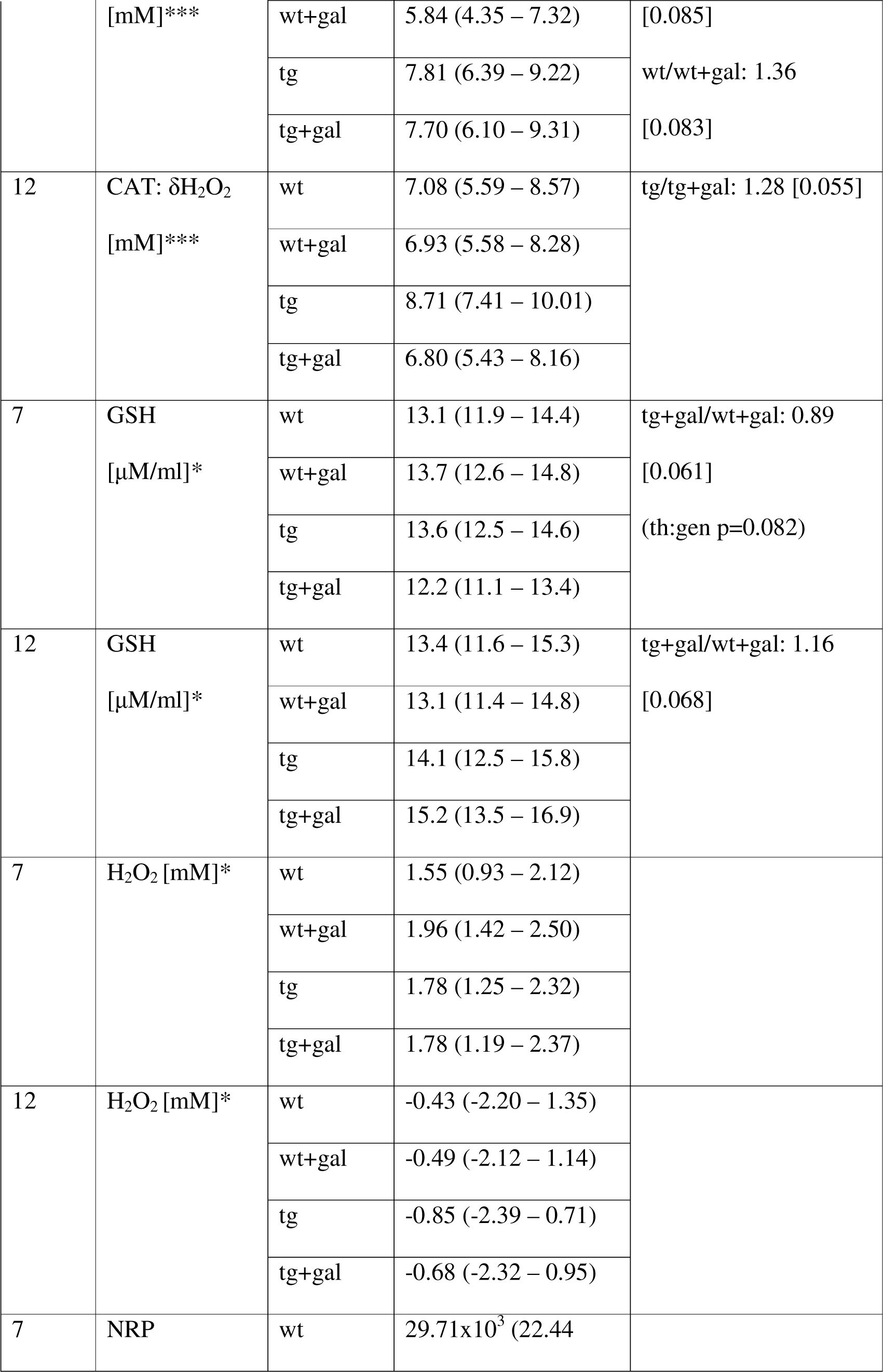
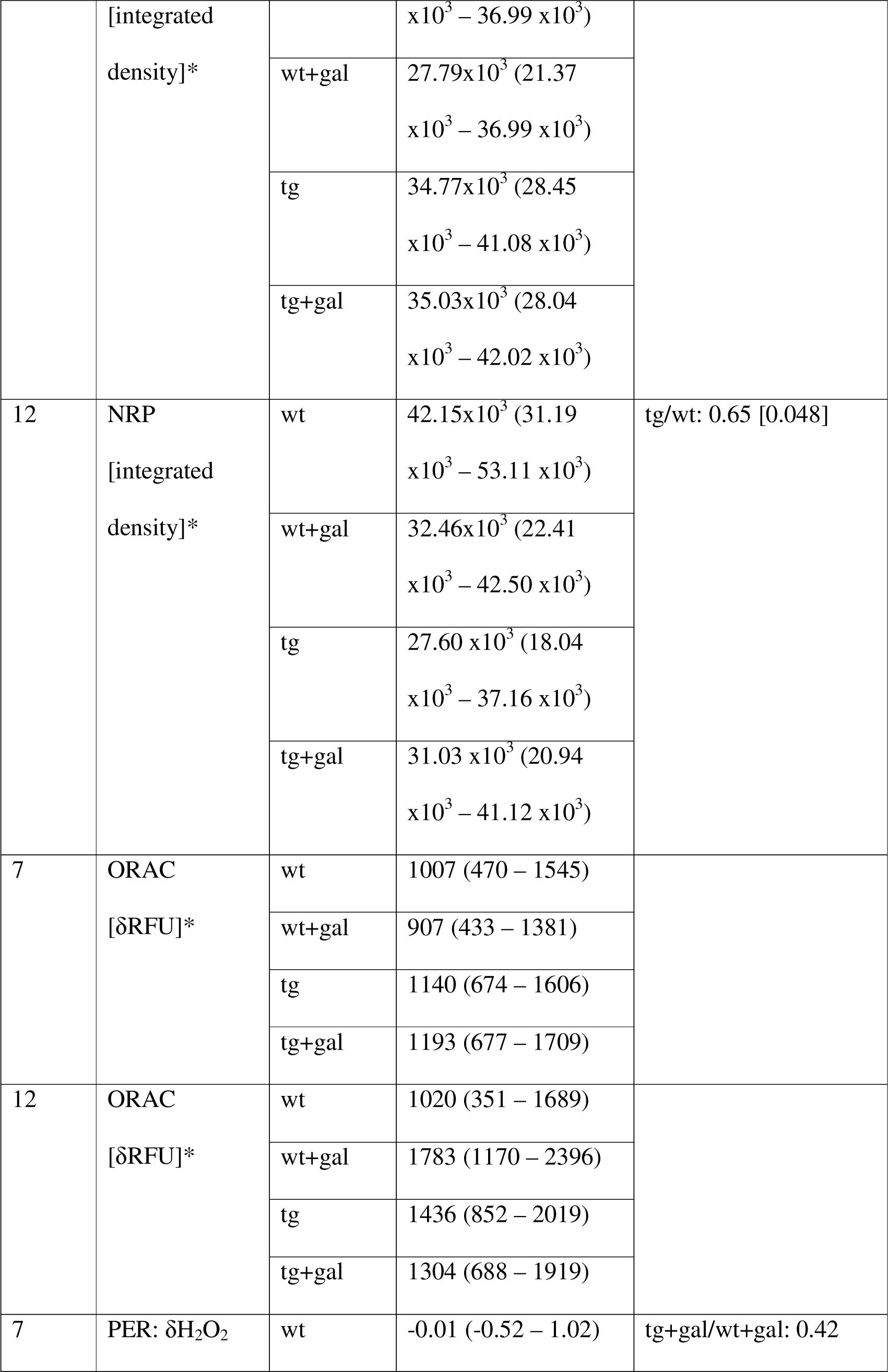
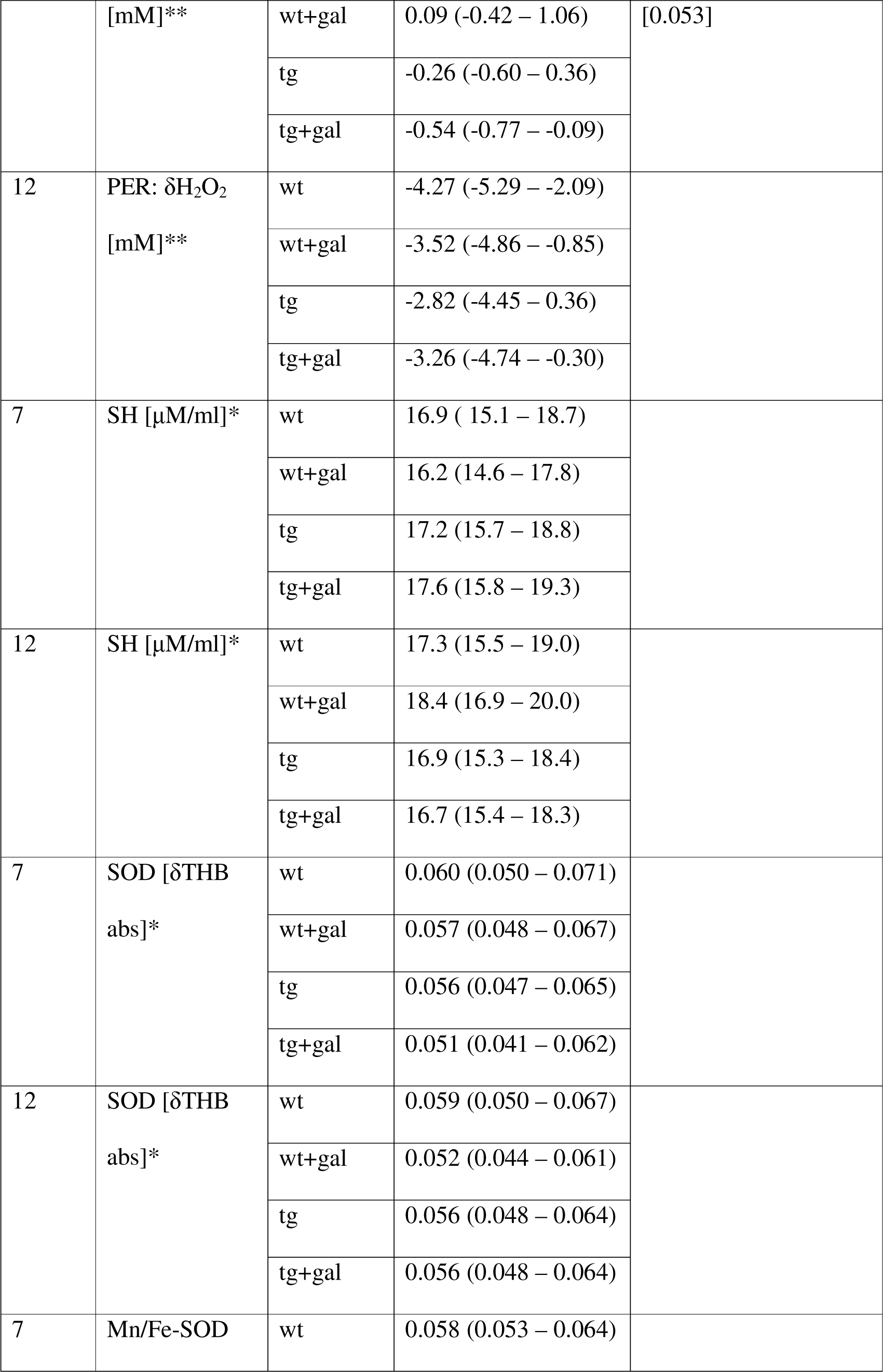
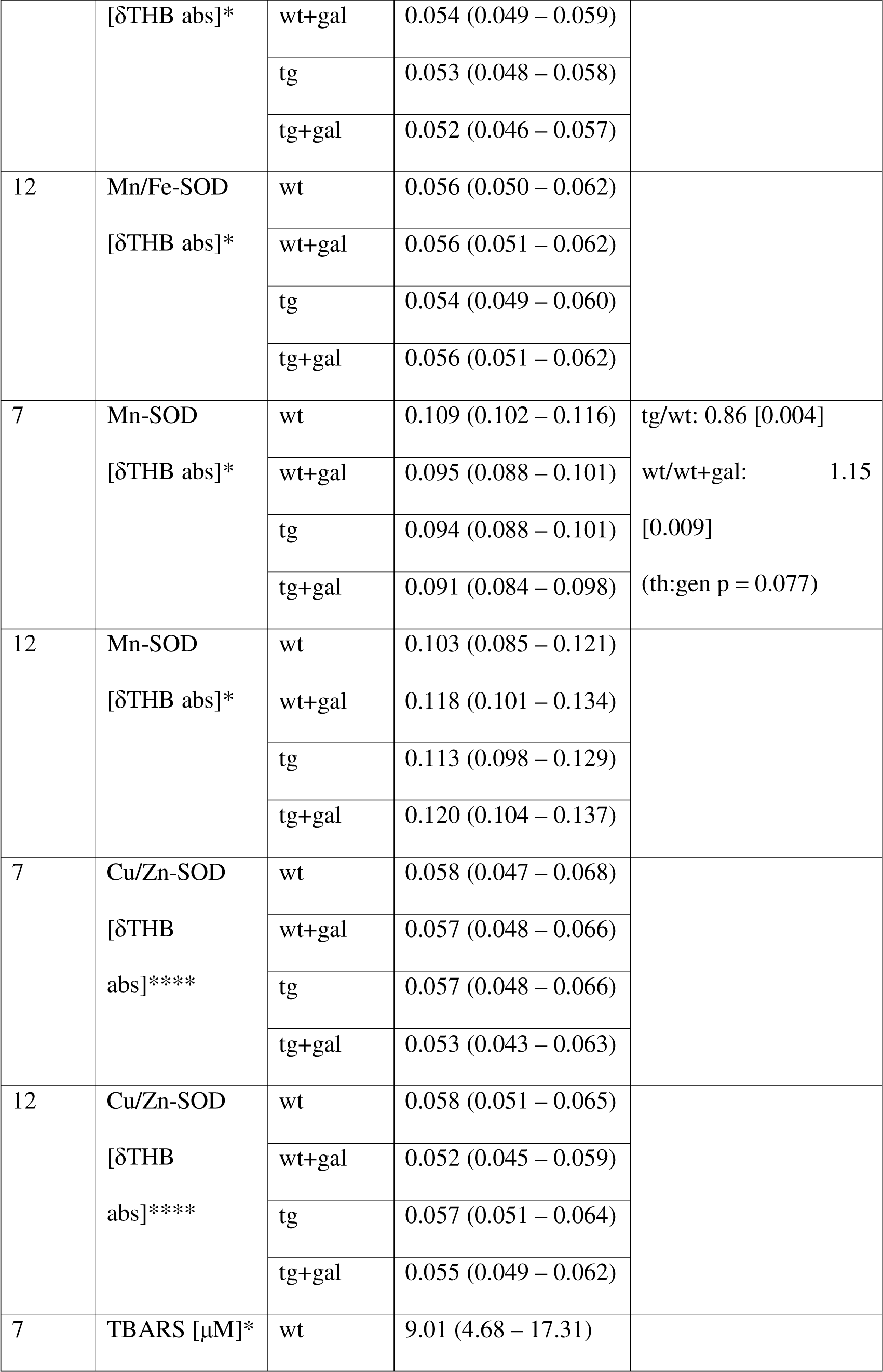
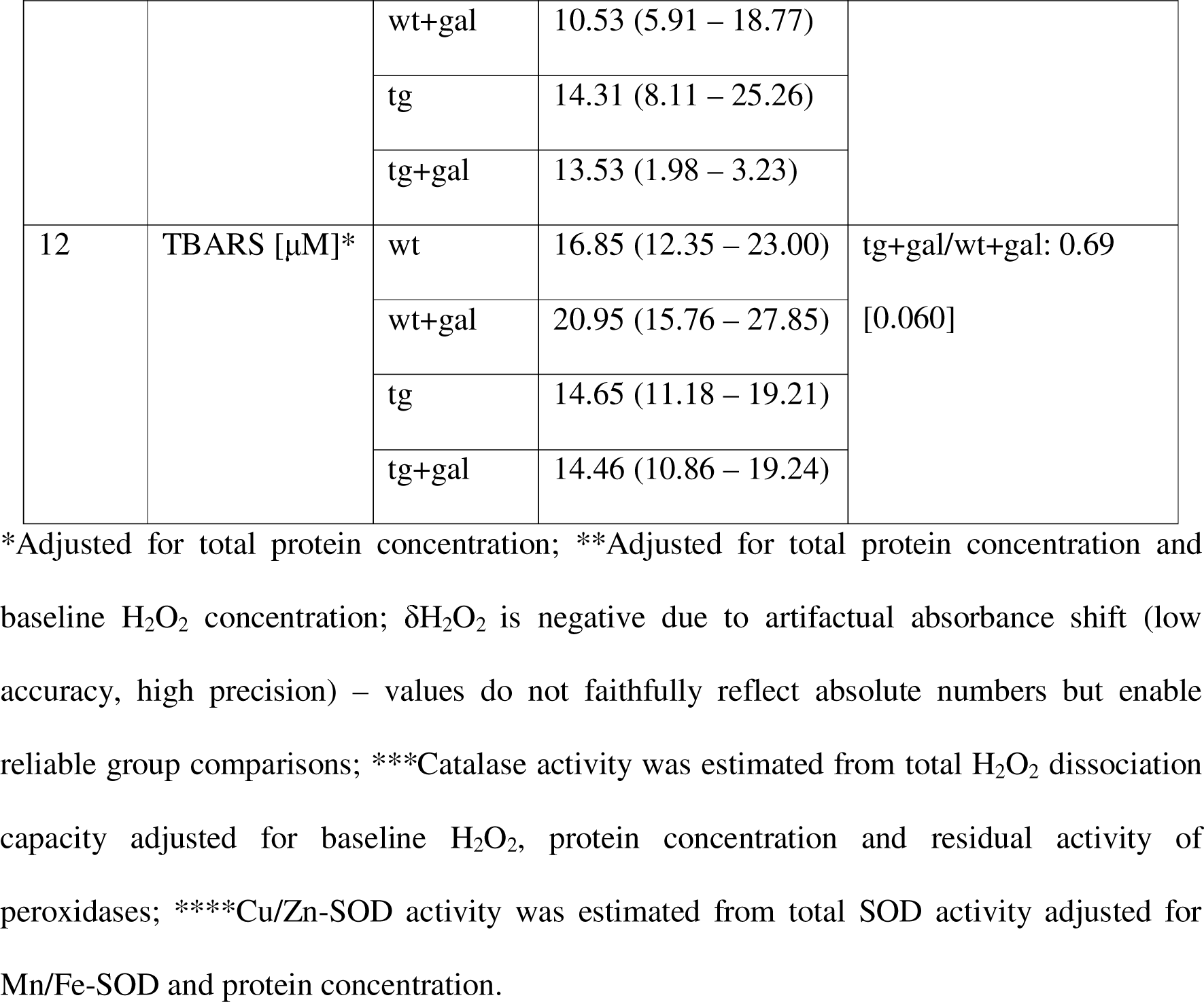
Gastrointestinal redox biomarkers in the presymptomatic (7-month-old) and symptomatic (12-month-old) Tg2576 mice.

In the symptomatic (12-month-old) Tg2576, total antioxidant capacity was reduced (NRP: - 35%; ORAC: +41% (inversely related to antioxidant capacity)) (Table 1; Figure 1). Total H_2_O_2_ dissociation capacity (+21%) and the activity of catalase (+23%) and peroxidases were increased, and there was a slight decrement in Mn-SOD capacity (+10% δTHB absorbance). However, the observed changes were associated with decreased accumulation of lipid peroxidation end products in the transgenic gastrointestinal tract (−13%) (Table 1; Figure 1). D-galactose treatment increased antioxidant capacity (NRP: +12%; ORAC: -9%) and decreased H_2_O_2_ dissociation capacity only in Tg2576 (−22%), possibly due to suppression of catalase activity (−22%). The observed change in H_2_O_2_ metabolism following D-galactose in the symptomatic Tg2576 was associated with a slight increment in GSH availability (+8%). Conversely, D-galactose treatment decreased antioxidant capacity in the gastrointestinal tract of wild-type animals (NRP: -23%; ORAC: +75%). Diminished antioxidant capacity was associated with increased activity of total (−12% δTHB absorbance) and cytoplasmic Cu/Zn-SOD (−11% δTHB absorbance) and decreased activity of mitochondrial Mn-SOD (+15% δTHB absorbance). The observed changes were accompanied by a +24% increased accumulation of lipid peroxidation end products in the wild-type animals receiving D-galactose (Table 1; Figure 1).

Of the observed changes, only gut antioxidant capacity (ABTS) was an important predictor of cognitive performance in the presymptomatic stage (Figure 1C). Mice with greater gut antioxidant capacity had better cognitive performance (Figure 1C). In the advanced stage, gastrointestinal antioxidant capacity (ABTS, NRP, ORAC), SOD capacity (total and Mn/Fe-SOD), low-molecular and protein thiols, and lipid peroxidation end products were determined to be important predictors of spatial memory (Figure 1C). Greater antioxidant capacity was associated with better cognitive performance based on NRP (r=-0.53) and ORAC (r=0.48), while ABTS assay (r=0.48) suggested an inverse association. Both total SOD (δTHB; r=- 0.52) and Mn/Fe-SOD (δTHB; r=-0.38) activity were associated with poor cognitive performance, while mitochondrial (Mn-SOD) and cytoplasmic (Cu/Zn-SOD) fractions did not predict spatial memory (Figure 1C). Interestingly, low-molecular-weight and protein thiols were both important predictors of cognitive performance; however, low-molecular-weight thiols were associated with poor spatial memory, while the inverse was true for protein thiol residues. Cognitive impairment was inversely proportional to the concentration of lipid peroxidation end products in the gastrointestinal tract.

Presymptomatic Tg2576 demonstrated increased exploration drive and decreased grooming and nesting (**Figure 2**). D-galactose treatment normalized exploration and grooming but did not increase the nesting score in the AD model. Conversely, there was no effect of D-galactose on measured behavioral parameters in the controls. Interestingly, there were no pronounced differences in exploration, grooming, or nesting in the cognitively deficient 12- month-old Tg2576, suggesting the observed phenotypic traits are specific to the presymptomatic phase. D-galactose treatment was associated with reduced grooming behavior and velocity in the OF test; however, only in the 12-month-old Tg2576 (**Figure 2**).

**Figure 2.**
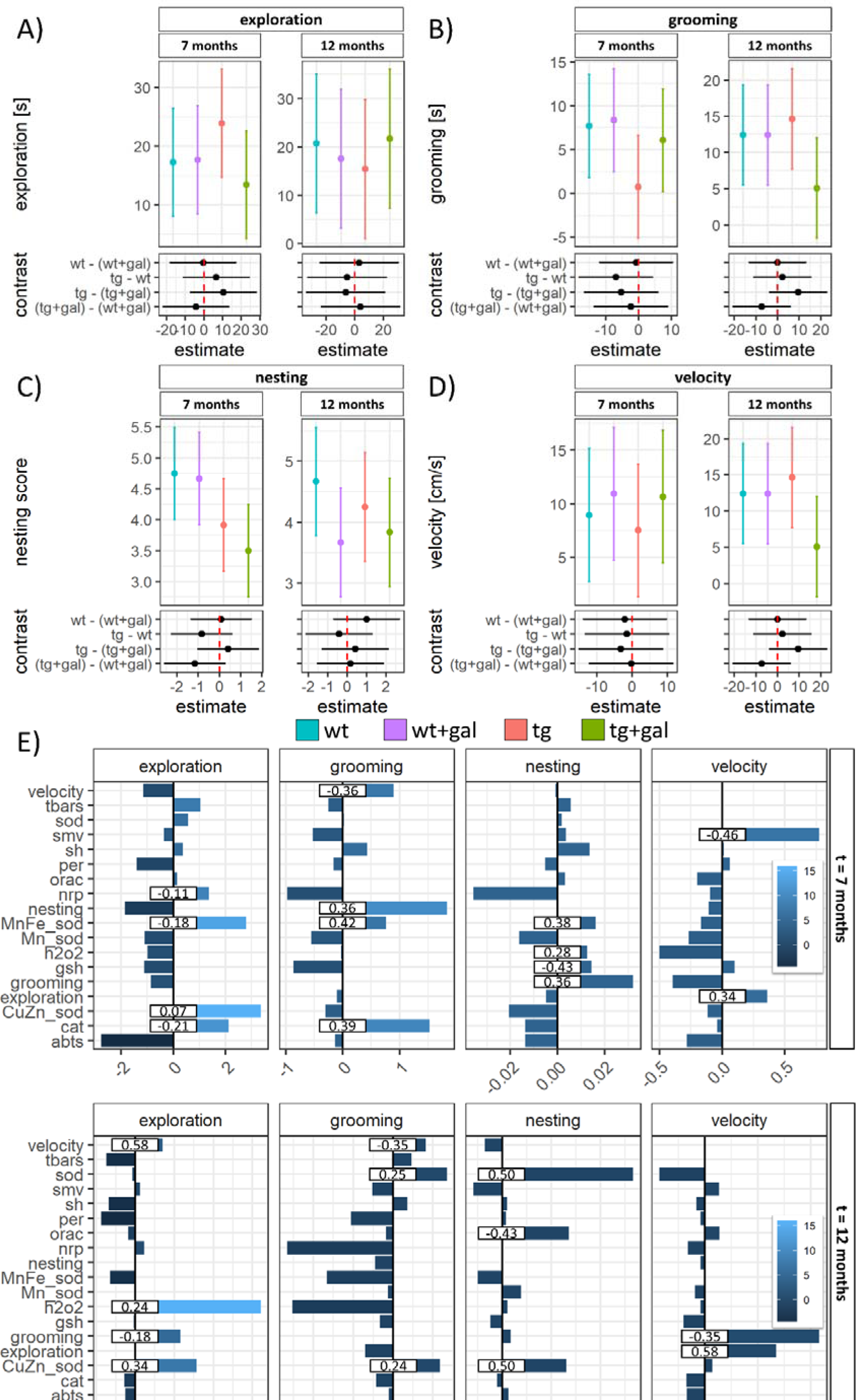
Behavioural characteristics of the presymptomatic (7-month-old) and symptomatic (12-month-old) Tg2576 mice after chronic oral D-galactose treatment (200 mg/kg). Model output depicting exploratory activity **(A)**, grooming **(B)**, nesting **(C)**, and velocity **(D)** point estimates and 95% confidence intervals (CI) in the presymptomatic (left) and symptomatic (right) Tg2576 mice and respective wild type controls treated with vehicle (wt; tg) or oral D-galactose (wt+gal; tg+gal). Group differences are presented as contrasts (point estimates and 95% CI of differences between group least square means). **E)** Variable importance maps calculated based on the permutation-induced mean decrease in accuracy derived from conditional inference-based unbiased classification random forests with exploration, grooming, nesting, and velocity defined as response variables. Bars indicate permutation importance for cforest models and numbers indicate correlation coefficients. abts – difference in absorbance of 2,2′-azino-bis(3-ethylbenzothiazoline-6-sulfonic acid); cat – catalase activity; Cu/Zn-sod – the activity of cytoplasmic Cu/Zn-superoxide dismutase; dh2o2 – total H2O2 dissociation capacity; gsh – glutathione; h2o2 – H2O2; Mn-sod – Mn-superoxide dismutase; Mn/Fe-sod – Mn- and Fe-superoxide dismutase; nrp – nitrocellulose redox permanganometry (antioxidant capacity biomarker); orac – oxygen radical absorbance capacity; per – residual activity of peroxidases; sh – protein thiol residues; sod – total superoxide dismutase activity; tbars – thiobarbituric acid reactive substances.

Mn/Fe-SOD was an important predictor of several phenotypic traits in presymptomatic animals (**Figure 2E**). Greater Mn/Fe-SOD activity was associated with exploration (δTHB; r=-0.18), and reduced grooming (δTHB; r=0.42) and nesting (δTHB; r=0.38) scores. The increased exploratory drive was also associated with decreased gut antioxidant capacity (NRP) and decreased gastrointestinal catalase activity (r=-0.21). Catalase activity was an important predictor of grooming (r=0.39) while nesting was associated with decreased low-molecular-weight thiols (r=-0.43) and increased concentration of H_2_O_2_ (r=0.28) **(Figure 2E)**.

In the symptomatic stage, increased H_2_O_2_ (r=0.24) and decreased Cu/Zn-SOD activity (δTHB; r=0.34) predicted exploratory drive. Total and cytoplasmic SOD activity was inversely associated with both grooming (δTHB; total: r=0.25; Cu/Zn-SOD: r=0.24) and nesting (δTHB; total: r=0.50; Cu/Zn-SOD: r=0.50). Nesting was also associated with greater gut antioxidant capacity (ORAC; r=-0.43) **(Figure 2E)**.

It has been hypothesized that redox biomarkers might be associated with GI function, and thus, possibly reflected in fecal pellet output. Total fecal pellet output was unremarkable both in the 7-month-old and the 12-month-old animals (**Figure 3A**). In the presymptomatic stage, chronic oral D-galactose treatment was associated with fewer pellets produced in the 7- month-old wild-type animals; however, there was no change in the Tg2576. There was no association between fecal pellet output and spatial memory in the 7-month-old mice (smv vs. fecal pellet output; r = 0.03); however, 12-month-old mice with poor spatial memory produced fewer pellets on average (smv vs. fecal pellet output; r = -0.32) (Figure 3B). Redox biomarkers demonstrated variable associations with fecal pellet output. The strongest association was observed for cytoplasmic SOD in the 7-month-old mice (Figure 3C) and protein thiol groups in the 12-month-old animals (Figure 3D). In both cases, greater antioxidant capacity was associated with the production of fewer pellets. Associations between fecal pellet output with all redox biomarkers are provided in Supplementary Material.

**Figure 3.**
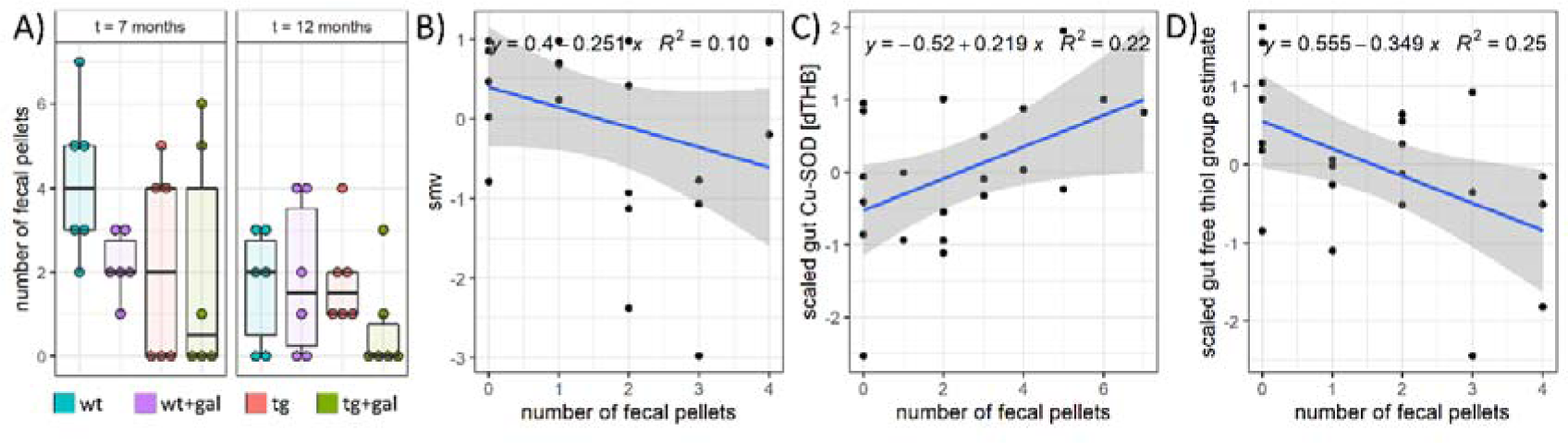
Fecal output is partially associated with redox parameters in the gut, but not with smv. Fecal output (A) and the association between fecal output and smv (B), scaled gut Cu-SOD (C), and protein sulfhydryls (D). smv – spatial memory vector; Cu-SOD – Cu/Zn superoxide dismutase.

Finally, we analysed the association between GI redox biomarkers measured in this study with the expression of plasma and brain metabolic and neuropathological markers measured by Babic Perhoc et al. in the same cohort of Tg2576 mice ^32^. Correlations between all parameters are provided in Supplementary Material.

## Discussion

Presented results suggest an increased electrophilic burden in the Tg2576 gut compensated by the time the animals develop cognitive deficits. Chronic oral D-galactose administration is associated with detrimental effects in the gut of wild-type animals but not in the Tg2576 GI tract providing evidence in favor of the hypothesis the beneficial effects of D-galactose might depend on the underlying pathophysiology. Gut redox biomarkers seem to be associated with behavioral patterns across groups with the most pronounced contribution related to spatial memory in the advanced stage of the disease possibly reflecting the behavioral effects mediated by the gut-brain axis^33^.

Although the effects were small and accompanied by large uncertainty (due to the exploratory nature of the experiment), redox biomarkers suggest an increased GI electrophilic burden might precede the development of cognitive deficits in Tg2576. The activity of mitochondrial antioxidant enzyme Mn-SOD was increased in the gut of presymptomatic Tg2576, possibly indicating a compensatory response to increased generation of O_2_^•-^ anions (Table 1, Figure 1,2). Increased expression and activity of Mn-SOD represent important mechanisms by which cells defend against the accumulation of free radicals generated in the process of mitochondrial respiration^34^. The activity of cytoplasmic Cu/Zn-SOD and bacterial Fe-SOD did not change, indicating that the electrophilic burden in the Tg2576 gut might be primarily related to mitochondrial metabolism. The hypothesis that increased Mn-SOD represents a compensatory response is supported by the finding of increased accumulation of TBARS indicating either failure of upstream antioxidant defense systems in preventing initiation of lipid peroxidation; or the inability of terminating mechanisms to stop its propagation^35^. In the symptomatic Tg2576, Mn-SOD activity was decreased, possibly as a result of prolonged exposure to increased concentration of free radicals^36^. Moreover, total antioxidant capacity was decreased and the activities of H_2_O_2_-metabolizing systems (catalase and peroxidases) were increased (Table 1, Figure 1,2). Increased wasting of nucleophilic substrates (reflected in reduced antioxidant capacity) and upregulation of H_2_O_2_ metabolism were accompanied by neutralization of lipid peroxidation suggesting that compensatory mechanisms were able to stabilize increased generation of free radicals by redefining a redox heterostatic setpoint most likely at the expense of long term functioning^18,37^. The observed results, suggesting an increased electrophilic burden in the presymptomatic stage heterostatically compensated by the time the animals developed cognitive deficits, are in line with previous reports in other animal models of familial AD. Chi et al. reported reduced expression of enzymes involved in GSH-mediated defense against oxidative stress and decreased total antioxidant capacity in the GI tract of the presymptomatic APP/PS1 mice^3^. Honarpisheh et al. did not assess GI redox homeostasis; however, they observed multiple pathophysiological alterations in the presymptomatic Tg2576 that did not persist to the symptomatic stage^4^. Early pathophysiological alterations observed by Honarpisheh et al. in the presymptomatic Tg2576 might be associated with unopposed electrophilic burden, while the establishment of redox heterostasis corresponds with the absence of structural and functional changes of the GI barrier in the aged mice^4^.

Chronic oral D-galactose treatment was associated with detrimental effects on redox homeostasis in the 7- and 12-month-old wild-type mice; however, in the Tg2576, the effects of D-galactose were either absent (presymptomatic stage) or associated with beneficial modulation of redox homeostasis (symptomatic stage). Chronic parenteral administration of D-galactose is widely utilized for the induction of oxidative stress and modeling aging ^38,39^ so its detrimental effects on redox homeostasis in wild-type animals are not surprising. Nevertheless, harmful effects of chronic D-galactose treatment are usually observed after parenteral administration of large doses, while chronic oral *ad libitum* administration has so far only been associated with beneficial effects in animal models of AD^25,26,32^ and atopic dermatitis^40^. A large body of evidence suggests that the majority of harmful and beneficial effects of D-galactose might be explained by tissue exposure. Peroral administration of D-galactose, associated with beneficial effects, usually results in 10-fold lower tissue exposure in comparison with parenteral routes due to intestinal absorption and liver retention^27,28,41,42^. In contrast, bolus oral administration can exceed tissue buffering capacity and achieve high plasma D-galactose concentrations^25^ and has, thus, been associated with detrimental effects^43,44^. Likewise, parenteral (i.e. subcutaneous) administration of D-galactose can exert beneficial effects^45^ if tissue metabolic capacity is not exceeded. Nevertheless, there are some exceptions (e.g. a recent report of beneficial effects of large-dose parenteral D-galactose in irradiated mice^46^) and it cannot be excluded that the effects of D-galactose are not only dose-but also model-dependent^41,47^. The presented results support this concept as detrimental effects of chronic oral D-galactose on gut redox homeostasis (and cognition) were observed only in the wild-type controls.

One of our aims was to explore whether GI redox biomarkers were valid predictors of mouse behavior. Recent evidence strongly supports the role of gut microbiota in regulating animal behavior^29–31^; however, exact mechanisms used by gut microbiota to control behavioral patterns are still not fully understood. Considering a close relationship between gut eubiosis and GI redox homeostasis^48–52^, gut redox milieu might be associated with specific behavioral patterns. In this research, GI antioxidant capacity (ABTS) was recognized as a potential predictor of cognitive performance across groups, with greater antioxidant capacity associated with better spatial memory in the 7-month-old mice. In the 12-month-old mice, several biomarkers were recognized as potential predictors of cognitive performance. NRP, ORAC, and concentration of protein thiols suggested that increased antioxidant capacity predicted better overall spatial memory; however, lipid peroxidation, GSH, and lower activity of total and Mn/Fe-SOD were also paradoxically recognized as positive predictors of cognitive performance. Discrepant results suggest the relationship between GI redox homeostasis and cognitive performance cannot be described by a simple model in which „better redox homeostasis“ translates to improved spatial memory. The understudied direct and indirect (e.g. blood flow-mediated) effects of meal ingestion on intraluminal redox homeostasis provide one possible explanation for the paradoxical findings. The latter is supported by the recognition of Mn/Fe-SOD as one of the predictors as this enzyme is found in plants and bacteria. In this context, reduced activity of Mn/Fe-SOD, associated with better spatial memory, might reflect either suppressed microbiota-derived SOD or the absence of food ingestion shortly before the trial. Unfortunately, temporal patterns of food intake were not monitored in the present study. Mn/Fe-SOD also predicted other behavioral patterns – its activity was associated with exploration and inversely associated with grooming and nesting scores in the 7-month-old cohort. Nesting was associated with greater baseline H_2_O_2_ and reduced GSH, while catalase activity acted as a positive predictor of grooming and a negative predictor of exploration. In the 12-month-old mice, total and cytoplasmic SOD activity demonstrated an inverse correlation with grooming and nesting, while the decreased activity of Cu/Zn-SOD and decreased baseline H_2_O_2_ predicted greater exploration. The latter might reflect inflammatory signaling associated with the generation of O_2_^•-^ ^53^ as it has been shown that even mild inflammation is sufficient to induce anxiety-like behavior^54^. More research is needed to understand the biological implications of the observed associations.

Finally, we found no evidence of a strong association between fecal pellet output and spatial memory. In the pre-symptomatic stage, there was no association, while in the symptomatic stage, there was only a weak association between cognitive ability and the number of fecal pellets. The number of fecal pellets was associated with some redox biomarkers possibly indicating the reduced antioxidant capacity in animals producing more pellets; however, the relationship between redox homeostasis and GI function remains to be elucidated.

## Conclusion

The presented results indicate an electrophilic challenge of the GI redox homeostasis in the Tg2576 mice before the development of cognitive deficits compensated by the time the animals develop symptoms. The results suggest that gut oxidative stress might be involved in the processes driving the initiation of neurodegenerative changes. Alternatively, subtle CNS changes taking place in the early stage of neurodegeneration might affect gut redox homeostasis while the animals are still not manifesting cognitive deficits. Future research should elucidate whether the observed changes might contribute to neurodegenerative processes by supporting peripheral inflammation.

### Limitations

The most important limitation of this study is its exploratory design with only 5-6 animals per group. Consequently, the power to detect changes in some redox biomarkers with satisfactory certainty was fairly low. For example, in the presymptomatic Tg2576 lipid peroxidation in the gut was increased by ∼60% - indicating a relatively large and likely biologically meaningful effect. Nevertheless, the effect did not meet the predetermined criteria for statistical significance due to a small number of animals and relatively large dispersion. We fixed α at 10% to minimize the risk of omission and avoid an excessive false nondiscovery rate; however, as evident from the abovementioned example, it is reasonable to assume that some relevant effects might have still been missed and additional studies are needed to better understand the role of GI redox homeostasis in transgenic models of AD. Another limitation is that temporal patterns of food intake were not monitored in the present study. As it can be speculated that the presence of intraluminal nutrients might exert both direct and indirect effects on redox homeostasis of the gut it cannot be ruled out that some of the effects reflected the difference in meal timing caused by either genotype or D-galactose treatment.

## Materials and methods

### Animals

Forty adult male B6;SJL-Tg(APPSWE)2576Kha heterozygous transgenic mice (Tg2576) and wild-type controls aged 5 months and 40 mice aged 10 months (Taconic Biosciences Inc., Hudson NY, USA) entered the experiments. Mice were housed individually in standard cages in vented positive pressure cabinets maintaining a stable temperature (22-24°C) and humidity (40-60%) environment, with a 12-h light/12-h dark cycle, at the licensed animal facility at the Croatian Institute for Brain Research, University of Zagreb School of Medicine (HR-POK-006). Mice were kept on standardized food pellets and water *ad libitum* before galactose treatment was initiated, after which treated groups received galactose dissolved in tap water in the previously observed daily water intake volume with tap water made available after the daily dose was drunk.

### Experimental design

The first mice cohort of 40 animals entered the experiment at 5-months-old, to assess the characteristics of presymptomatic fAD, whereas the second cohort investigating symptomatic fAD started the experiment at 10-months-old. In both instances, mice were divided into 4 groups with 10 animals per group; Tg2576 (tg), wild types (wt), galactose-treated Tg2576 (tg+gal) and galactose-treated wild types (wt+gal). Oral D-galactose treatment lasted for 2 months, when the animals underwent cognitive and behavioural testing as indicated below (aged 7 and 12 months, respectively). Following these tests, animals were sacrificed and samples withdrawn for further analyses. Tissue samples used in this study were from the same animals used in the study by Babic Perhoc et al. for the assessment of the effects of D-galactose on metabolic parameters in the CNS ^32^.

### Galactose treatment

Oral D-galactose treatment lasted for two months, and was initiated in the two cohorts at 5 and 10 months of age, respectively. D-galactose was freshly dissolved in tap water at a concentration of 200 mg/kg/day, in a volume previously observed as average daily water intake for each animal (∼10 ml daily). Control Tg2576 and wild-type mice received regular tap water throughout the experiment.

### Behavioral analysis

Behavioral assessment was based on a battery of tests including the Morris Water Maze (MWM), Open Field (OF), and nesting assessment. Technical details of MWM and OF are described in detail in the original publication ^32^. Spatial memory parameters from the original study ^32^ were dedimenzionalized into a single spatial memory vector parameter (described in Data Analysis). Grooming, velocity, and exploration drive were determined in the OF. Nesting was performed according to a pre-defined protocol ^55^ before sacrification. Individually housed mice were given a small rectangle (approximately 5 x 5 cm) of pressed cotton an hour before the dark cycle. The following morning, each cage was assessed by two observers and given a score on a rating scale of 1-5, as indicated in ^55^, with 1 representing the lowest score and more than 90% of the piece of cotton left untouched and 5 representing a near perfect nest. The average score from both investigators was recorded for each built nest.

### Fecal pellet output

Fecal pellet output was measured by counting the number of fecal pellets produced in the OF arena during 300 s.

### Sample preparation

The animals were euthanized by decapitation in deep anesthesia (thiopental 60 mg/kg/diazepam 6 mg/kg ip). Internal organs were exposed by laparotomy and the duodenum was removed and snap-frozen in liquid nitrogen and stored at -80°C. Frozen tissue sections were thawed in lysis buffer (150 mM NaCl, 50 mM Tris-HCl pH 7.4, 1 mM EDTA, 1% Triton X-100, 1% sodium deoxycholate, 0.1% SDS, 1 mM PMSF, protease (Sigma-Aldrich, Burlington, MA, USA) and phosphatase (PhosSTOP, Roche, Basel, Switzerland) inhibitor cocktail; pH 7.5) and subjected to 3 cycles of ultrasonic homogenization (Microson Ultrasonic Cell 167 Disruptor XL, Misonix, Farmingdale, NY, SAD). Homogenates were centrifuged for 10 min (relative centrifugal force 12 879 g) and protein concentration was measured using the Bradford reagent (Sigma-Aldrich, USA). A protein calibration curve was determined with bovine serum albumin dissolved in a lysis buffer. Homogenates were kept at -80 °C.

### Antioxidant capacity

Antioxidant capacity was determined with 3 separate biochemical methods. The 2,2′-azino-bis(3-ethylbenzothiazoline-6-sulfonic acid) (ABTS) radical cation assay was performed by measuring the change in absorbance of ABTS working solution after incubation in the presence of tissue samples in a 96 microwell plate. ABTS (7 mM) was incubated with K_2_S_2_O_8_ (2.45 mM) for 24 hours. ABTS/K_2_S_2_O_8_ solution was diluted 40-fold and incubated with 1 μl of each homogenate in a volumetric ratio of 1:100 (sample/ABTS). The absorbance at 405 nm was measured after 5 min using an Infinite F200 PRO multimodal microplate reader (Tecan, Switzerland) ^10^. The difference between the baseline control solution (ABTS) and final absorbance was proportional to the sample antioxidant capacity ^56^. The oxygen radical absorbance capacity (ORAC) fluorescein assay was conducted by incubating tissue samples (10 μl) with 150 μl of 5 μM fluorescein dissolved in phosphate-buffered saline (pH 7) in a 96 microwell plate ^57^. Fluorescence (465 nm excitation/540 nm emission; 10 nm bandwidth) was recorded every minute for 60 min using an Infinite F200 PRO multimodal microplate reader (Tecan, Switzerland). Nitrocellulose redox permanganometry (NRP) was done by quantifying sample-mediated reduction of KMnO_4_ by measuring the integrated density of MnO_2_ precipitate on a nitrocellulose membrane. Briefly, 1 μl of each sample was pipetted onto the nitrocellulose membrane (Amersham Protran 0.45; GE Healthcare Life Sciences, Chicago, IL, USA), and the dry membrane was incubated in NRP solution (0.01 g/ml KMnO_4_ in ddH_2_O) for 30 s. The membrane was destained in ddH_2_O, digitalized, and analyzed in Fiji with the gel analyzer plugin ^58^.

### Catalase/peroxidase activity and H_2_O_2_ concentration

The activity of catalase and peroxidases and baseline concentration of H_2_O_2_ were measured by modified Hadwan’s assay ^59^ as described in ^60,61^. Method validation has been reported in^62^. Briefly, tissue samples (5 μl) were incubated with 150 μl of the Co(NO_3_)_2_ solution (0.1 g of Co(NO_3_)_2_×6 H_2_O in 5 ml ddH_2_O mixed with (NaPO_3_)_6_ solution (0.05 g (NaPO_3_)_6_ dissolved in 5 ml ddH_2_O) and added to 90 ml of NaHCO_3_ solution (8.1 g in 90 ml ddH_2_O)) in a microwell plate to obtain baseline H_2_O_2_ concentrations. The same procedure was repeated after incubating the samples with 10 mM H_2_O_2_ in PBS for 90 seconds to measure total H_2_O_2_ dissociation capacity and in the presence of 10 mM H_2_O_2_ and 0.025 mM NaN_3_ in PBS to measure NaN_3_-insensitive fraction reflecting the activity of peroxidases ^61,63^. The H_2_O_2_ concentration was estimated from the ([Co(CO_3_)_3_]Co) absorbance at 450 nm using the Infinite F200 PRO multimodal microplate reader (Tecan, Switzerland) based on the linear model obtained with serial dilutions of H_2_O_2_ in PBS (R^2^ = 0.97 – 0.99). Assay sensitivity was optimized by modifying sample volumes and reaction times based on pilot experiments on the same samples.

### SOD activity

The activity of superoxide dismutase (SOD) was analysed utilizing an indirect assay based on the inhibition of 1,2,3-trihydroxybenzene (THB) autoxidation introduced by Marklund and Marklund ^64^ and modified by others ^28,65^. Briefly, tissue samples (6 μl) were incubated with 100 μl of the SOD working solution (THB solution (80 μL; 60 mM THB in 1 mM HCl) mixed with 4000 μL of the reaction buffer (0.05 M Tris-HCl and 1 mM Na_2_EDTA in ddH_2_O; pH 8.2) in a 96-well plate for 300 s. Absorbance increment at 450 nm reflecting THB autoxidation was measured with kinetic intervals of 30 s to approximate SOD activity. The same procedure was repeated with modified reaction buffers containing inhibitors partially selective for Cu/Zn-SOD (2 mM KCN) or Cu/Zn-SOD and Fe-SOD (5 mM H_2_O_2_) ^61,63^.

### Protein and low-molecular-weight thiol group quantification

Free thiol groups were quantified with Ellman’s procedure based on the reaction of sulfhydryl residues with 5,5’-dithio-bis-(2-nitrobenzoic acid)(DTNB) which yields a yellow-colored product 5-thio-2-nitrobenzoic acid (TNB)^9,27,66^. Tissue homogenates (25 μl) were mixed with an equal amount of 4% sulfosalicylic acid solution for 60 min on ice and the samples were centrifuged for 10 min at 10 000 g to obtain the protein (pellet) and low-molecular-weight (supernatant) fractions. Both fractions were incubated with the DTNB solution (4 mg/ml DTNB in 5% sodium citrate) at room temperature for 10 min and 405 nm absorbance was measured with the Infinite F200 PRO multimodal microplate reader (Tecan, Switzerland). The concentration of thiol residues was estimated based on the extinction coefficient of 14 150 M^-1^cm^-1^.

### Lipid peroxidation

Lipid peroxidation was measured with the modified thiobarbituric acid reactive substances (TBARS) assay ^67^ as described previously ^27,28^. Briefly, tissue homogenates (12 μl) were incubated with 120 μl of the TBA-TCA reagent (0.4% thiobarbituric acid in 15% trichloroacetic acid) in a heating block set at 95 °C for 20 min in perforated microcentrifuge tubes. The coloured adduct was extracted with n-butanol (100 μl) and the absorbance was measured at 540 nm using the Infinite F200 PRO multimodal microplate reader (Tecan, Switzerland). The concentration of TBARS was determined from the linear model obtained with serial dilutions of malondialdehyde (Sigma-Aldrich, USA)(R^2^ = 0.98 – 0.99). The procedure was optimized for the analysis of duodenal homogenates by iterative adjustment of sample volume, reaction time, and the n-butanol extraction procedure.

### Data analysis

Data were analysed using R (4.1.3) following ARRIVE 2.0 guidelines for reporting animal studies ^68^. Gastrointestinal redox biomarkers were analysed by fitting linear models using the biomarker of interest as the dependent variable, while group allocation (based on the treatment and genotype) and protein concentration (loading control) were defined as independent predictors. Additional covariates were introduced where necessary: baseline concentration of H_2_O_2_ (total H_2_O_2_ dissociation capacity, catalase, and peroxidase activity models); peroxidase activity (catalase activity model); Mn/Fe-SOD activity (Cu/Zn-SOD activity model). The effect of age was not modelled directly as it was not permitted by the experimental design (separate cohorts). A change in THB absorbance (inversely proportional to SOD activity) was used in modelling instead of approximated activity measures of SOD to reduce artifactual uncertainty. Visual inspection of residuals was performed to check model assumptions and appropriate (log) transformations were introduced where appropriate. Model outputs (group least square means and their contrasts) were reported as point estimates with 95% confidence intervals (CI). Contrasts were reported as ratios regardless of the presence of log transformation in the original model to facilitate the interpretation of effect sizes. Considering the exploratory nature of the study, α was set at 10% to deflate the type II error and avoid excessive false nondiscovery rate (particularly for large effects) (e.g. the threshold for -log_10_(p) was fixed at 1 in the volcano plot) ^69^. Complex variables (e.g. MWM indices of spatial memory) were de-dimensionalized with principal component analysis (conducted on a centered, scaled, and mean-imputed dataset) to obtain a single biologically meaningful variable (e.g. spatial memory vector (SMV)). SMV for the presymptomatic and symptomatic stages captured 66.2% and 94.7% of the variance (quadrant preference in training and test trials), respectively. Correlation matrices were based on Pearson and Spearman correlation coefficients. Variable importance was determined using the permutation-induced mean decrease in accuracy derived from conditional inference-based unbiased classification random forests in the computational toolbox for recursive partitioning (party)^70^.

## Author contributions

ABP, AK, JOB – treatment, behavioural experiments; JH, ABP – biochemical analyses, data curation, data analysis, writing the first draft of the manuscript. AK, JOB, DV, MSP – critical revision of the manuscript. MSP – funding, supervision.

## Data Availability

Raw data can be obtained from the corresponding author. The manuscript has been preprinted on bioRxiv (https://doi.org/10.1101/2023.06.03.542513).

## Ethics approval

The animal procedures were conducted in concordance with current institutional (University of Zagreb School of Medicine), national (The Animal Protection Act, NN135/2006; NN 47/2011), and international (Directive 2010/63/EU) guidelines on the use of experimental animals. The experiments were approved by the Croatian Ministry of Agriculture (EP 186/2018; 525-10/0255-15-5) and the Ethical Committee of the University of Zagreb School of Medicine (380-59-10106-18-111/173).

## Consent to participate

Not applicable.

## Consent for publication

Not applicable.

## Acknowledgments

None.

## References

(1) Singh, A.; Dawson, T. M.; Kulkarni, S. Neurodegenerative Disorders and Gut-Brain Interactions. J Clin Invest 2021, 131 (13), e143775. https://doi.org/10.1172/JCI143775.

(2) Sun, M.; Ma, K.; Wen, J.; Wang, G.; Zhang, C.; Li, Q.; Bao, X.; Wang, H. A Review of the Brain-Gut-Microbiome Axis and the Potential Role of Microbiota in Alzheimer’s Disease. J Alzheimers Dis 2020, 73 (3), 849–865. https://doi.org/10.3233/JAD-190872.

(3) Chi, H.; Cao, W.; Zhang, M.; Su, D.; Yang, H.; Li, Z.; Li, C.; She, X.; Wang, K.; Gao, X.; Ma, K.; Zheng, P.; Li, X.; Cui, B. Environmental Noise Stress Disturbs Commensal Microbiota Homeostasis and Induces Oxi-Inflammmation and AD-like Neuropathology through Epithelial Barrier Disruption in the EOAD Mouse Model. J Neuroinflammation 2021, 18 (1), 9. https://doi.org/10.1186/s12974-020-02053-3.

(4) Honarpisheh, P.; Reynolds, C. R.; Blasco Conesa, M. P.; Moruno Manchon, J. F.; Putluri, N.; Bhattacharjee, M. B.; Urayama, A.; McCullough, L. D.; Ganesh, B. P. Dysregulated Gut Homeostasis Observed Prior to the Accumulation of the Brain Amyloid-β in Tg2576 Mice. Int J Mol Sci 2020, 21 (5), 1711. https://doi.org/10.3390/ijms21051711.

(5) Manocha, G. D.; Floden, A. M.; Miller, N. M.; Smith, A. J.; Nagamoto-Combs, K.; Saito, T.; Saido, T. C.; Combs, C. K. Temporal Progression of Alzheimer’s Disease in Brains and Intestines of Transgenic Mice. Neurobiol Aging 2019, 81, 166–176. https://doi.org/10.1016/j.neurobiolaging.2019.05.025.

(6) Semar, S.; Klotz, M.; Letiembre, M.; Van Ginneken, C.; Braun, A.; Jost, V.; Bischof, M.; Lammers, W. J.; Liu, Y.; Fassbender, K.; Wyss-Coray, T.; Kirchhoff, F.; Schäfer, K.-H. Changes of the Enteric Nervous System in Amyloid-β Protein Precursor Transgenic Mice Correlate with Disease Progression. J Alzheimers Dis 2013, 36 (1), 7–20. https://doi.org/10.3233/JAD-120511.

(7) Wang, Y.; An, Y.; Ma, W.; Yu, H.; Lu, Y.; Zhang, X.; Wang, Y.; Liu, W.; Wang, T.; Xiao, R. 27- Hydroxycholesterol Contributes to Cognitive Deficits in APP/PS1 Transgenic Mice through Microbiota Dysbiosis and Intestinal Barrier Dysfunction. J Neuroinflammation 2020, 17 (1), 199. https://doi.org/10.1186/s12974-020-01873-7.

(8) Homolak, J.; Babic Perhoc, A.; Knezovic, A.; Osmanovic Barilar, J.; Koc, F.; Stanton, C.; Ross, R. P.; Salkovic-Petrisic, M. Disbalance of the Duodenal Epithelial Cell Turnover and Apoptosis Accompanies Insensitivity of Intestinal Redox Homeostasis to Inhibition of the Brain Glucose-Dependent Insulinotropic Polypeptide Receptors in a Rat Model of Sporadic Alzheimer’s Disease. Neuroendocrinology 2022, 112 (8), 744–762. https://doi.org/10.1159/000519988.

(9) Homolak, J.; Babic Perhoc, A.; Knezovic, A.; Osmanovic Barilar, J.; Salkovic-Petrisic, M. Failure of the Brain Glucagon-Like Peptide-1-Mediated Control of Intestinal Redox Homeostasis in a Rat Model of Sporadic Alzheimer’s Disease. Antioxidants (Basel) 2021, 10 (7), 1118. https://doi.org/10.3390/antiox10071118.

(10) Homolak, J.; De Busscher, J.; Zambrano-Lucio, M.; Joja, M.; Virag, D.; Babic Perhoc, A.; Knezovic, A.; Osmanovic Barilar, J.; Salkovic-Petrisic, M. Altered Secretion, Constitution, and Functional Properties of the Gastrointestinal Mucus in a Rat Model of Sporadic Alzheimer’s Disease. ACS Chem Neurosci 2023. https://doi.org/10.1021/acschemneuro.3c00223.

(11) Qian, X.-H.; Liu, X.-L.; Chen, G.; Chen, S.; Tang, H.-D. Injection of Amyloid-β to Lateral Ventricle Induces Gut Microbiota Dysbiosis in Association with Inhibition of Cholinergic Anti-Inflammatory Pathways in Alzheimer’s Disease. J Neuroinflammation 2022, 19 (1), 236. https://doi.org/10.1186/s12974-022-02599-4.

(12) Alves, S. S.; Silva-Junior, R. M. P. da; Servilha-Menezes, G.; Homolak, J.; Šalković-Petrišić, M.; Garcia-Cairasco, N. Insulin Resistance as a Common Link Between Current Alzheimer’s Disease Hypotheses. J Alzheimers Dis 2021, 82 (1), 71–105. https://doi.org/10.3233/JAD-210234.

(13) Brown, G. C. The Endotoxin Hypothesis of Neurodegeneration. J Neuroinflammation 2019, 16 (1), 180. https://doi.org/10.1186/s12974-019-1564-7.

(14) Homolak, J. Targeting the Microbiota-Mitochondria Crosstalk in Neurodegeneration with Senotherapeutics. Adv Protein Chem Struct Biol 2023, 136, 339–383. https://doi.org/10.1016/bs.apcsb.2023.02.018.

(15) Qian, X.-H.; Song, X.-X.; Liu, X.-L.; Chen, S.; Tang, H.-D. Inflammatory Pathways in Alzheimer’s Disease Mediated by Gut Microbiota. Ageing Res Rev 2021, 68, 101317. https://doi.org/10.1016/j.arr.2021.101317.

(16) Soto, M.; Herzog, C.; Pacheco, J. A.; Fujisaka, S.; Bullock, K.; Clish, C. B.; Kahn, C. R. Gut Microbiota Modulate Neurobehavior through Changes in Brain Insulin Sensitivity and Metabolism. Mol Psychiatry 2018, 23 (12), 2287–2301. https://doi.org/10.1038/s41380-018-0086-5.

(17) Butterfield, D. A.; Halliwell, B. Oxidative Stress, Dysfunctional Glucose Metabolism and Alzheimer Disease. Nat Rev Neurosci 2019, 20 (3), 148–160. https://doi.org/10.1038/s41583-019-0132-6.

(18) Homolak, J. Redox Homeostasis in Alzheimer’s Disease. In Redox Signaling and Biomarkers in Ageing; Çakatay, U., Ed.; Healthy Ageing and Longevity; Springer International Publishing: Cham, 2022; pp 323–348. https://doi.org/10.1007/978-3-030-84965-8_15.

(19) Vaccaro, A.; Kaplan Dor, Y.; Nambara, K.; Pollina, E. A.; Lin, C.; Greenberg, M. E.; Rogulja, D. Sleep Loss Can Cause Death through Accumulation of Reactive Oxygen Species in the Gut. Cell 2020, 181 (6), 1307–1328.e15. https://doi.org/10.1016/j.cell.2020.04.049.

(20) McCord, J. M. Radical Explanations for Old Observations. Gastroenterology 1987, 92 (6), 2026– 2028. https://doi.org/10.1016/0016-5085(87)90640-8.

(21) Homolak, J. Gastrointestinal Redox Homeostasis in Ageing. Biogerontology 2023.

(22) Bischoff, S. C.; Barbara, G.; Buurman, W.; Ockhuizen, T.; Schulzke, J.-D.; Serino, M.; Tilg, H.; Watson, A.; Wells, J. M. Intestinal Permeability--a New Target for Disease Prevention and Therapy. BMC Gastroenterol 2014, 14, 189. https://doi.org/10.1186/s12876-014-0189-7.

(23) Bhattacharyya, A.; Chattopadhyay, R.; Mitra, S.; Crowe, S. E. Oxidative Stress: An Essential Factor in the Pathogenesis of Gastrointestinal Mucosal Diseases. Physiol Rev 2014, 94 (2), 329– 354. https://doi.org/10.1152/physrev.00040.2012.

(24) Circu, M. L.; Aw, T. Y. Intestinal Redox Biology and Oxidative Stress. Semin Cell Dev Biol 2012, 23 (7), 729–737. https://doi.org/10.1016/j.semcdb.2012.03.014.

(25) Knezovic, A.; Osmanovic Barilar, J.; Babic, A.; Bagaric, R.; Farkas, V.; Riederer, P.; Salkovic-Petrisic, M. Glucagon-like Peptide-1 Mediates Effects of Oral Galactose in Streptozotocin-Induced Rat Model of Sporadic Alzheimer’s Disease. Neuropharmacology 2018, 135, 48–62. https://doi.org/10.1016/j.neuropharm.2018.02.027.

(26) Salkovic-Petrisic, M.; Osmanovic-Barilar, J.; Knezovic, A.; Hoyer, S.; Mosetter, K.; Reutter, W. Long-Term Oral Galactose Treatment Prevents Cognitive Deficits in Male Wistar Rats Treated Intracerebroventricularly with Streptozotocin. Neuropharmacology 2014, 77, 68–80. https://doi.org/10.1016/j.neuropharm.2013.09.002.

(27) Homolak, J.; Babic Perhoc, A.; Knezovic, A.; Kodvanj, I.; Virag, D.; Osmanovic Barilar, J.; Riederer, P.; Salkovic-Petrisic, M. Is Galactose a Hormetic Sugar? An Exploratory Study of the Rat Hippocampal Redox Regulatory Network. Mol Nutr Food Res 2021, 65 (21), e2100400. https://doi.org/10.1002/mnfr.202100400.

(28) Homolak, J.; Babic Perhoc, A.; Knezovic, A.; Osmanovic Barilar, J.; Virag, D.; Joja, M.; Salkovic-Petrisic, M. The Effect of Acute Oral Galactose Administration on the Redox System of the Rat Small Intestine. Antioxidants (Basel) 2021, 11 (1), 37. https://doi.org/10.3390/antiox11010037.

(29) Hsiao, E. Y.; McBride, S. W.; Hsien, S.; Sharon, G.; Hyde, E. R.; McCue, T.; Codelli, J. A.; Chow, J.; Reisman, S. E.; Petrosino, J. F.; Patterson, P. H.; Mazmanian, S. K. Microbiota Modulate Behavioral and Physiological Abnormalities Associated with Neurodevelopmental Disorders. Cell 2013, 155 (7), 1451–1463. https://doi.org/10.1016/j.cell.2013.11.024.

(30) Luczynski, P.; McVey Neufeld, K.-A.; Oriach, C. S.; Clarke, G.; Dinan, T. G.; Cryan, J. F. Growing up in a Bubble: Using Germ-Free Animals to Assess the Influence of the Gut Microbiota on Brain and Behavior. Int J Neuropsychopharmacol 2016, 19 (8), pyw020. https://doi.org/10.1093/ijnp/pyw020.

(31) Needham, B. D.; Funabashi, M.; Adame, M. D.; Wang, Z.; Boktor, J. C.; Haney, J.; Wu, W.-L.; Rabut, C.; Ladinsky, M. S.; Hwang, S.-J.; Guo, Y.; Zhu, Q.; Griffiths, J. A.; Knight, R.; Bjorkman, P. J.; Shapiro, M. G.; Geschwind, D. H.; Holschneider, D. P.; Fischbach, M. A.; Mazmanian, S. K. A Gut-Derived Metabolite Alters Brain Activity and Anxiety Behaviour in Mice. Nature 2022, 602 (7898), 647–653. https://doi.org/10.1038/s41586-022-04396-8.

(32) Babic Perhoc, A.; Osmanovic Barilar, J.; Knezovic, A.; Farkas, V.; Bagaric, R.; Svarc, A.; Grünblatt, E.; Riederer, P.; Salkovic-Petrisic, M. Cognitive, Behavioral and Metabolic Effects of Oral Galactose Treatment in the Transgenic Tg2576 Mice. Neuropharmacology 2019, 148, 50–67. https://doi.org/10.1016/j.neuropharm.2018.12.018.

(33) Carabotti, M.; Scirocco, A.; Maselli, M. A.; Severi, C. The Gut-Brain Axis: Interactions between Enteric Microbiota, Central and Enteric Nervous Systems. Ann Gastroenterol 2015, 28 (2), 203– 209.

(34) Candas, D.; Li, J. J. MnSOD in Oxidative Stress Response-Potential Regulation via Mitochondrial Protein Influx. Antioxid Redox Signal 2014, 20 (10), 1599–1617. https://doi.org/10.1089/ars.2013.5305.

(35) Ayala, A.; Muñoz, M. F.; Argüelles, S. Lipid Peroxidation: Production, Metabolism, and Signaling Mechanisms of Malondialdehyde and 4-Hydroxy-2-Nonenal. Oxid Med Cell Longev 2014, 2014, 360438. https://doi.org/10.1155/2014/360438.

(36) Demicheli, V.; Quijano, C.; Alvarez, B.; Radi, R. Inactivation and Nitration of Human Superoxide Dismutase (SOD) by Fluxes of Nitric Oxide and Superoxide. Free Radic Biol Med 2007, 42 (9), 1359–1368. https://doi.org/10.1016/j.freeradbiomed.2007.01.034.

(37) Ursini, F.; Maiorino, M.; Forman, H. J. Redox Homeostasis: The Golden Mean of Healthy Living. Redox Biol 2016, 8, 205–215. https://doi.org/10.1016/j.redox.2016.01.010.

(38) Sadigh-Eteghad, S.; Majdi, A.; McCann, S. K.; Mahmoudi, J.; Vafaee, M. S.; Macleod, M. R. D-Galactose-Induced Brain Ageing Model: A Systematic Review and Meta-Analysis on Cognitive Outcomes and Oxidative Stress Indices. PLoS One 2017, 12 (8), e0184122. https://doi.org/10.1371/journal.pone.0184122.

(39) Shwe, T.; Pratchayasakul, W.; Chattipakorn, N.; Chattipakorn, S. C. Role of D-Galactose-Induced Brain Aging and Its Potential Used for Therapeutic Interventions. Exp Gerontol 2018, 101, 13– 36. https://doi.org/10.1016/j.exger.2017.10.029.

(40) Kim, D.-Y.; Jung, D.-H.; Song, E.-J.; Jang, A.-R.; Park, J.-Y.; Ahn, J.-H.; Lee, T.-S.; Kim, Y.-J.; Lee, Y.- J.; Seo, I.-S.; Kim, H.-E.; Ryu, E.-J.; Sim, J.; Park, J.-H. D-Galactose Intake Alleviates Atopic Dermatitis in Mice by Modulating Intestinal Microbiota. Front Nutr 2022, 9, 895837. https://doi.org/10.3389/fnut.2022.895837.

(41) Homolak, J.; Babic Perhoc, A.; Virag, D.; Knezovic, A.; Osmanovic Barilar, J.; Salkovic-Petrisic, M. D-Galactose Might Protect against Ionizing Radiation by Stimulating Oxidative Metabolism and Modulating Redox Homeostasis. J Radiat Res 2023, 64 (4), 743–745. https://doi.org/10.1093/jrr/rrad046.

(42) Homolak, J.; Babić Perhoč, A.; Virag, D.; Knezovic, A.; Osmanovic, J.; Salkovic-Petrisic, M. D-Galactose Might Mediate Some of the Skeletal Muscle Hypertrophy-Promoting Effects of Milk - a Nutrient to Consider for Sarcopenia? Preprint 2023. https://doi.org/10.13140/RG.2.2.25134.59205.

(43) Budni, J.; Pacheco, R.; da Silva, S.; Garcez, M. L.; Mina, F.; Bellettini-Santos, T.; de Medeiros, J.; Voss, B. C.; Steckert, A. V.; Valvassori, S. da S.; Quevedo, J. Oral Administration of D-Galactose Induces Cognitive Impairments and Oxidative Damage in Rats. Behav Brain Res 2016, 302, 35– 43. https://doi.org/10.1016/j.bbr.2015.12.041.

(44) Budni, J.; Garcez, M. L.; Mina, F.; Bellettini-Santos, T.; da Silva, S.; Luz, A. P. da; Schiavo, G. L.; Batista-Silva, H.; Scaini, G.; Streck, E. L.; Quevedo, J. The Oral Administration of D-Galactose Induces Abnormalities within the Mitochondrial Respiratory Chain in the Brain of Rats. Metab Brain Dis 2017, 32 (3), 811–817. https://doi.org/10.1007/s11011-017-9972-9.

(45) Chogtu, B.; Arivazhahan, A.; Kunder, S. K.; Tilak, A.; Sori, R.; Tripathy, A. Evaluation of Acute and Chronic Effects of D-Galactose on Memory and Learning in Wistar Rats. Clin Psychopharmacol Neurosci 2018, 16 (2), 153–160. https://doi.org/10.9758/cpn.2018.16.2.153.

(46) Zhu, T.; Wang, Z.; He, J.; Zhang, X.; Zhu, C.; Zhang, S.; Li, Y.; Fan, S. D-Galactose Protects the Intestine from Ionizing Radiation-Induced Injury by Altering the Gut Microbiome. J Radiat Res 2022, 63 (6), 805–816. https://doi.org/10.1093/jrr/rrac059.

(47) Salkovic-Petrisic, M. Oral Galactose Provides a Different Approach to Incretin-Based Therapy of Alzheimer’s Disease. Journal of Neurology & Neuromedicine 2018, 3 (4).

(48) Circu, M. L.; Aw, T. Y. Redox Biology of the Intestine. Free Radic Res 2011, 45 (11–12), 1245– 1266. https://doi.org/10.3109/10715762.2011.611509.

(49) Jones, R. M.; Neish, A. S. Redox Signaling Mediated by the Gut Microbiota. Free Radic Biol Med 2017, 105, 41–47. https://doi.org/10.1016/j.freeradbiomed.2016.10.495.

(50) Million, M.; Tidjani Alou, M.; Khelaifia, S.; Bachar, D.; Lagier, J.-C.; Dione, N.; Brah, S.; Hugon, P.; Lombard, V.; Armougom, F.; Fromonot, J.; Robert, C.; Michelle, C.; Diallo, A.; Fabre, A.; Guieu, R.; Sokhna, C.; Henrissat, B.; Parola, P.; Raoult, D. Increased Gut Redox and Depletion of Anaerobic and Methanogenic Prokaryotes in Severe Acute Malnutrition. Sci Rep 2016, 6, 26051. https://doi.org/10.1038/srep26051.

(51) Pérez, S.; Taléns-Visconti, R.; Rius-Pérez, S.; Finamor, I.; Sastre, J. Redox Signaling in the Gastrointestinal Tract. Free Radic Biol Med 2017, 104, 75–103. https://doi.org/10.1016/j.freeradbiomed.2016.12.048.

(52) Reese, A. T.; Cho, E. H.; Klitzman, B.; Nichols, S. P.; Wisniewski, N. A.; Villa, M. M.; Durand, H. K.; Jiang, S.; Midani, F. S.; Nimmagadda, S. N.; O’Connell, T. M.; Wright, J. P.; Deshusses, M. A.; David, L. A. Antibiotic-Induced Changes in the Microbiota Disrupt Redox Dynamics in the Gut. eLife 2018, 7, e35987. https://doi.org/10.7554/eLife.35987.

(53) Tian, T.; Wang, Z.; Zhang, J. Pathomechanisms of Oxidative Stress in Inflammatory Bowel Disease and Potential Antioxidant Therapies. Oxid Med Cell Longev 2017, 2017, 4535194. https://doi.org/10.1155/2017/4535194.

(54) Bercik, P.; Verdu, E. F.; Foster, J. A.; Macri, J.; Potter, M.; Huang, X.; Malinowski, P.; Jackson, W.; Blennerhassett, P.; Neufeld, K. A.; Lu, J.; Khan, W. I.; Corthesy-Theulaz, I.; Cherbut, C.; Bergonzelli, G. E.; Collins, S. M. Chronic Gastrointestinal Inflammation Induces Anxiety-like Behavior and Alters Central Nervous System Biochemistry in Mice. Gastroenterology 2010, 139 (6), 2102–2112.e1. https://doi.org/10.1053/j.gastro.2010.06.063.

(55) Deacon, R. M. J. Assessing Nest Building in Mice. Nat Protoc 2006, 1 (3), 1117–1119. https://doi.org/10.1038/nprot.2006.170.

(56) Ilyasov, I. R.; Beloborodov, V. L.; Selivanova, I. A.; Terekhov, R. P. ABTS/PP Decolorization Assay of Antioxidant Capacity Reaction Pathways. Int J Mol Sci 2020, 21 (3), 1131. https://doi.org/10.3390/ijms21031131.

(57) Dávalos, A.; Gómez-Cordovés, C.; Bartolomé, B. Extending Applicability of the Oxygen Radical Absorbance Capacity (ORAC-Fluorescein) Assay. J Agric Food Chem 2004, 52 (1), 48–54. https://doi.org/10.1021/jf0305231.

(58) Homolak, J.; Kodvanj, I.; Babic Perhoc, A.; Virag, D.; Knezovic, A.; Osmanovic Barilar, J.; Riederer, P.; Salkovic-Petrisic, M. Nitrocellulose Redox Permanganometry: A Simple Method for Reductive Capacity Assessment. MethodsX 2022, 9, 101611. https://doi.org/10.1016/j.mex.2021.101611.

(59) Hadwan, M. H. Simple Spectrophotometric Assay for Measuring Catalase Activity in Biological Tissues. BMC Biochem 2018, 19 (1), 7. https://doi.org/10.1186/s12858-018-0097-5.

(60) Homolak, J. In Vitro Analysis of Catalase and Superoxide Dismutase Mimetic Properties of Blue Tattoo Ink. Free Radic Res 2022, 56 (5–6), 343–357. https://doi.org/10.1080/10715762.2022.2102976.

(61) Homolak, J.; Joja, M.; Grabaric, G.; Schiatti, E.; Virag, D.; Perhoc, A. B.; Knezovic, A.; Barilar, J. O.; Salkovic-Petrisic, M. The Absence of Gastrointestinal Redox Dyshomeostasis in the Brain-First Rat Model of Parkinson’s Disease Induced by Bilateral Intrastriatal 6-Hydroxydopamine. bioRxiv 2022, 2022.08.22.504759. https://doi.org/10.1101/2022.08.22.504759.

(62) Homolak, J. The Effect of a Color Tattoo on the Local Skin Redox Regulatory Network: An N-of-1 Study. Free Radic Res 2021, 55 (3), 221–229. https://doi.org/10.1080/10715762.2021.1912340.

(63) Ma, X.; Deng, D.; Chen, W.; Ma, X.; Deng, D.; Chen, W. Inhibitors and Activators of SOD, GSH-Px, and CAT; IntechOpen, 2017. https://doi.org/10.5772/65936.

(64) Marklund, S.; Marklund, G. Involvement of the Superoxide Anion Radical in the Autoxidation of Pyrogallol and a Convenient Assay for Superoxide Dismutase. Eur J Biochem 1974, 47 (3), 469– 474. https://doi.org/10.1111/j.1432-1033.1974.tb03714.x.

(65) Li, X. Improved Pyrogallol Autoxidation Method: A Reliable and Cheap Superoxide-Scavenging Assay Suitable for All Antioxidants. J Agric Food Chem 2012, 60 (25), 6418–6424. https://doi.org/10.1021/jf204970r.

(66) Anderson, M. E. Determination of Glutathione and Glutathione Disulfide in Biological Samples. Methods Enzymol 1985, 113, 548–555. https://doi.org/10.1016/s0076-6879(85)13073-9.

(67) Aguilar Diaz De Leon, J.; Borges, C. R. Evaluation of Oxidative Stress in Biological Samples Using the Thiobarbituric Acid Reactive Substances Assay. J Vis Exp 2020, No. 159. https://doi.org/10.3791/61122.

(68) Percie du Sert, N.; Ahluwalia, A.; Alam, S.; Avey, M. T.; Baker, M.; Browne, W. J.; Clark, A.; Cuthill, I. C.; Dirnagl, U.; Emerson, M.; Garner, P.; Holgate, S. T.; Howells, D. W.; Hurst, V.; Karp, N. A.; Lazic, S. E.; Lidster, K.; MacCallum, C. J.; Macleod, M.; Pearl, E. J.; Petersen, O. H.; Rawle, F.; Reynolds, P.; Rooney, K.; Sena, E. S.; Silberberg, S. D.; Steckler, T.; Würbel, H. Reporting Animal Research: Explanation and Elaboration for the ARRIVE Guidelines 2.0. PLoS Biol 2020, 18 (7), e3000411. https://doi.org/10.1371/journal.pbio.3000411.

(69) Amrhein, V.; Korner-Nievergelt, F.; Roth, T. The Earth Is Flat (p > 0.05): Significance Thresholds and the Crisis of Unreplicable Research. PeerJ 2017, 5, e3544. https://doi.org/10.7717/peerj.3544.

(70) Strobl, C.; Boulesteix, A.-L.; Zeileis, A.; Hothorn, T. Bias in Random Forest Variable Importance Measures: Illustrations, Sources and a Solution. BMC Bioinformatics 2007, 8, 25. https://doi.org/10.1186/1471-2105-8-25.

