## Supplementary figures and images for "An exploratory study of gastrointestinal redox biomarkers in the presymptomatic and symptomatic Tg2576 mouse model of familial Alzheimer’s disease – phenotypic correlates and the effects of chronic oral D-galactose"

### Supplemental Fig 1

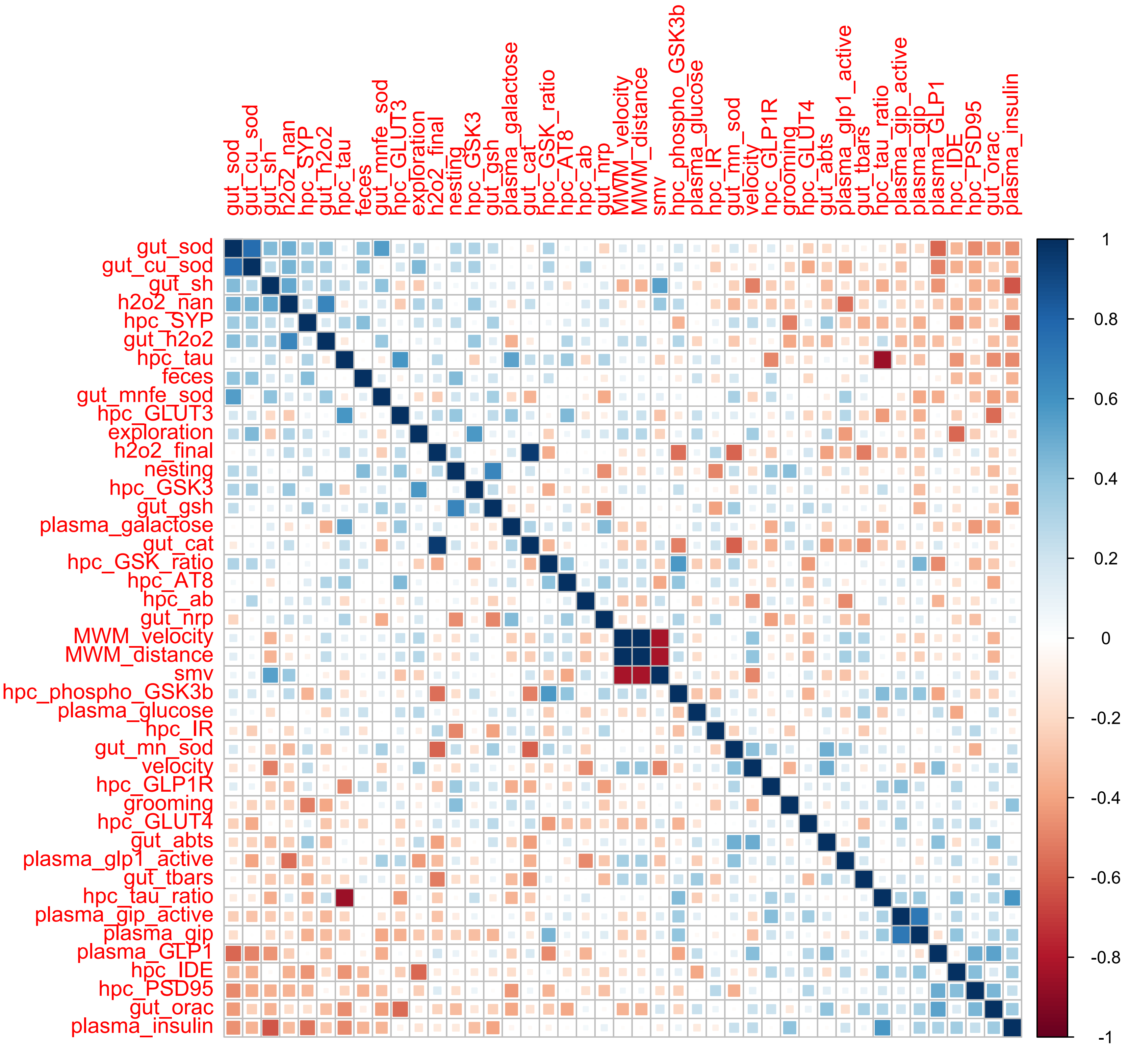

### Supplemental Fig 2

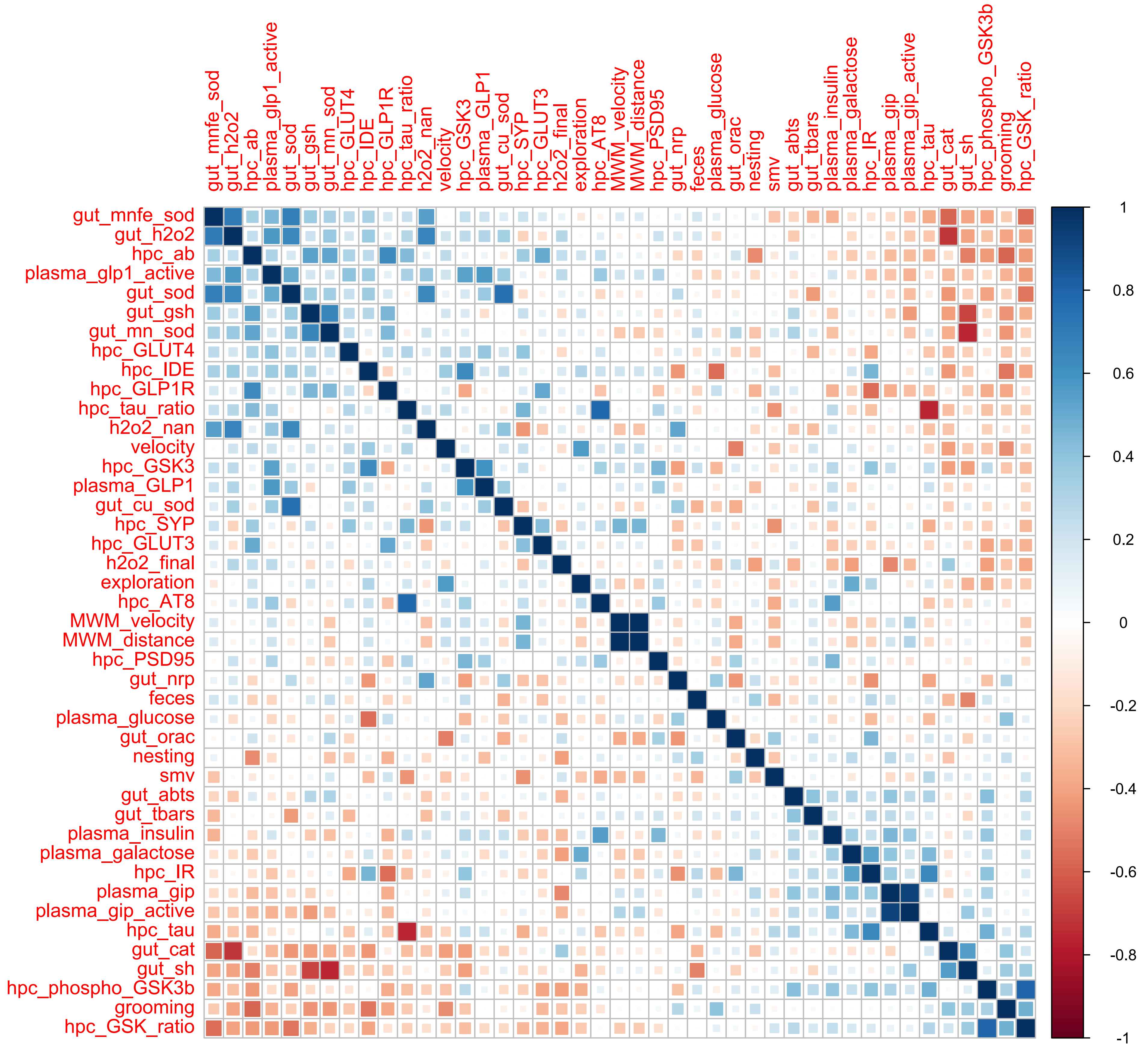
